# Genetic variation reveals individual-level climate tracking across the annual cycle of a migratory bird

**DOI:** 10.1101/2020.04.15.043331

**Authors:** Rachael A. Bay, Daniel S. Karp, James F. Saracco, William R.L. Anderegg, Luke O. Frishkoff, David Wiedenfeld, Thomas B. Smith, Kristen Ruegg

## Abstract

For migratory species, seasonal movements complicate local climate adaptation, as it is unclear whether individuals track climate niches across the annual cycle. In the migratory songbird yellow warbler (*Setophaga petechia*), we find a correlation between individual-level wintering and breeding precipitation but not temperature. Birds wintering in the driest regions of the Neotropics breed in the driest regions of North America. Individuals from drier regions also possess distinct morphologies and population responses to varying rainfall. We find a positive association between bill size and breeding season precipitation which, given documented climate-associated genomic variation, might reflect adaptation to local precipitation regimes. Relative abundance in the breeding range is linked to interannual fluctuations in precipitation, but the directionality of this response varies across geography. Together, our results suggest that variation in climate optima may exist across the breeding range of yellow warblers and provide a mechanism for selection across the annual cycle.

## Introduction

Intraspecific variation in climate-associated traits constitutes the raw material needed for species to adapt to ongoing climate change. Spatial environmental gradients can promote divergence in phenotypes and genotypes, providing a mechanism for the maintenance of variation in contemporary populations that could be beneficial under future climate regimes (Aitken *et al.* 2008; Bay & Palumbi 2014; Walters & Berger 2019). At the same time, narrow population-level climate niches may render populations maladapted to rapid environmental change (Brady *et al.* 2019). Outcomes under climate change will depend on the amount of standing genetic variation, the strength of selection pressure imposed by rapid environmental change, and the degree of gene flow among populations. While it is becoming increasingly clear that evolution will be required for some species to persist through the next century (Rehfeldt *et al.* 2002; Jump & Penuelas 2005), much remains to be understood about the distribution and maintenance of putatively adaptive variation across contemporary climate gradients.

Understanding links between genotype, phenotype, and climate is particularly challenging for migratory species, which face selective constraints on both their breeding and wintering grounds. Selection on wintering grounds could either reinforce or counteract selection on breeding grounds (Saino *et al.* 2004; Gunnarsson *et al.* 2005). At the species level, the question of whether environmental conditions are similar or different during different migratory stages has been investigated by assessing the role of climate in setting distributional limits of migratory species (Gómez *et al.* 2016; Zurell *et al.* 2018; Somveille *et al.* 2019). Some species, classified as ‘niche switchers,’ have been shown to migrate between areas with very different climates. Others, classified as ‘niche trackers,’ migrate between regions that share more similar climatic conditions than would be expected by chance. The biotic and abiotic environmental components of a species niche are extremely broad, however, much of the niche-tracking literature focuses on the particular role of climate in determining the ranges of migratory species (Gómez *et al.* 2016). Though the relative frequency of niche tracking versus switching across migratory species is still debated (Gómez *et al.* 2016; Somveille *et al.* 2019; Winger *et al.* 2019), niche tracking has been proposed as a mechanism explaining the evolution of migration, where species migrate to escape seasonality. In New World warblers (Parulidae), for example, migratory species tend to have more similar breeding and wintering climates than resident species (Gómez *et al.* 2016).

To date, analyses of potential climatic constraints on the seasonal distributions of migratory species have focused nearly entirely on the species level with little consideration of patterns within species. Could individuals also be tracking their climate niches such that, for example, individual birds from the hottest regions of the wintering range migrate to the hottest locations within the species’ breeding range? Previously, investigation of this question was limited by the requirement of range-wide, individual-level definitions of migratory connections between the breeding and wintering ranges. Stable isotope analysis, for example, provides individual-level information but not at high-enough resolution to link climatic conditions between an individual’s breeding and wintering grounds (Hobson 2005; Kelly *et al.* 2005). Like stable isotopes, geolocator studies provide individual-level information, but suffer from small sample sizes due to their cost and the fact that they require individuals to be recaptured after migrating (Bridge *et al.* 2013). Despite these limitations, such studies have provided a rich understanding individual movement in relationship to resource availability and abiotic conditions (Thorup *et al.* 2017; MacPherson *et al.* 2018). For example, a recent study on white storks (*Ciconia ciconia*) found evidence that individuals tracked climate across their migratory cycle. While that study represents a major advance in our understanding of within-species niche tracking, it was limited to individuals caught at a single location (Fandos *et al.* 2020). Genetic assignment has the potential to overcome issues of both resolution and sampling size, but these approaches have historically focused on the population or subspecies level (Ruegg *et al.* 2014). Fortunately, recent technological and analytical advances now make it possible to take advantage of isolation by distance signals in genetic data in order to estimate an individual’s natal site within the breeding range with high accuracy (Rañola *et al.* 2014).

We leverage such advances to ask whether there is evidence for individual-level climate tracking in the Neotropical migratory songbird yellow warbler (*Setophaga petechia*). On the species level, there is a high degree of overlap between breeding and wintering precipitation regimes across the range of yellow warblers, but less so for temperature (Gómez *et al.* 2016). Our previous work has shown that there is genetic variation associated with precipitation across the species’ breeding range and that mismatches between climate and genotype may have contributed to population declines over the last half century (Bay *et al.* 2018). Here, we use genomic data to describe migratory connectivity across the full annual cycle of the yellow warbler and test whether individuals track similar climates across their annual cycles. Because climate tracking is a potential means for maintaining local adaptation to climate across the annual cycle, we further test whether morphological traits show associations with climate and analyze how climate covaries with population size fluctuations over the past 50 years.

## Methods

### Sampling and Genotyping

We assembled 663 yellow warbler blood, tissue, and feather samples taken across the breeding and wintering ranges (Figure S1, Table S1). These samples constitute a collection from previous studies (Milot *et al.* 2000; Boulet *et al.* 2006), tissue from museum collections, and blood or feathers taken at banding stations. DNA was extracted from all samples using the Qiagen DNeasy Blood and Tissue Kit. Blood and tissue samples from the breeding range were used to create RAD-Seq libraries and the remainder of samples were genotyped using custom Fluidigm SNPtype assays as described below.

Single nucleotide polymorphisms (SNPs) from RAD-Seq data generated for Bay et al. (Bay *et al.* 2018) were used to examine population structure across the breeding range. Detailed methods can be found in that paper. Briefly, we used the bestRAD protocol (Ali *et al.* 2016) to create libraries for 229 breeding individuals, which were sequenced on an Illumina HiSeq 4000. Paired end 150bp reads were quality filtered using scripts from the STACKS pipeline (Catchen *et al.* 2013) and aligned to the yellow warbler genome (Bay *et al.* 2018) using bowtie2 (Langmead & Salzberg 2012). We called SNPs with GATK following best practices (www.broadinstitute.org). Using the R package genoscapeRtools, we discarded low coverage SNPs and low coverage individuals, resulting in a final dataset of 104,711 SNPs across 195 individuals. Within this dataset, all individuals had less than 25% missing data (mean=3.8%), each SNP had less than 10% missing genotypes (mean=3.8%), and we discarded SNPs with minor allele frequency less than 1%.

RAD-Seq data were used to select a panel of SNPs used to assign individuals to breeding groups. We used four different analyses: 1) *F*_*ST*_ between easternmost (Newfoundland, Nova Scotia, New Brunswick) and westernmost (British Columbia, Washington) locations, 2) *F*_*ST*_ between northernmost (Alaska and Churchill, Manitoba) and southernmost (Oregon, Pennsylvania, Michigan) locations, 3) Standard linear models comparing allele frequency to latitude, and 4) Standard linear models comparing allele frequency to longitude. Per locus *F*_*ST*_ was calculated using the Weir and Cockerham method in the R package hierfstat (Goudet 2005). For each of these metrics, we took the top ranked SNPs (highest *F*_*ST*_ or lowest p-value), for a total of 192 SNPs to be tested for genotyping. We developed Fluidigm SNPtype assays, which allow us to leverage the low yield DNA available in feather samples (Ruegg *et al.* 2014), for these 192 SNPs and used these assays to genotype 231 feather samples from 28 new locations across the breeding range. These data were used to determine a final set of 96 SNPs based on the quality of the genotyping assay and linear correlations with latitude and longitude. These 96 SNPs were genotyped in 203 birds from across the wintering range (Figure S1; Table S1). Only individuals with at least 80% of SNPs genotyped were used in analysis. A summary of the samples genotyped with each SNP set can be found in Table S2.

### Population structure and migratory connectivity

To define breeding populations, we ran STRUCTURE (Pritchard *et al.* 2000) on all high-quality SNPs from the 192 SNP panel, combining Fluidigm and RAD-Seq genotypes. We ran a range of K values (1-6) with 5 runs each (BURNIN=500000, NUMREPS=150000, LOCPRIOR=1) and used the Evanno method to identify the best-fit K value. Breeding populations were mapped spatially using the method implemented in TESS3 (Caye *et al.* 2016), to interpolate ancestry coefficients across the geographic range. Map locations were colored by the dominant ancestry cluster and transparency indicates the percent ancestry of that cluster, with the largest value assigned as opaque. We used population structure across the North American breeding range as a baseline to assign wintering birds to breeding populations. We used the R package rubias (Anderson *et al.* 2008) to first investigate how reliable our markers were to assign individuals to breeding populations using leave-one-out self-assignment. We then assigned each wintering sample back to a pre-defined breeding population (PofZ>0.8).

In addition to visualizing migratory connectivity based on discrete groups, we also used principal components analysis (PCA) using the R package SNPRelate (Zheng *et al.* 2012) to describe patterns on a continuous scale. We ran three separate PCAs for breeding range samples: 1) all SNPs in the RAD-Seq dataset, 2) SNPs genotyped for all breeding samples, and 3) the 96 SNPs used for assignment. Standard linear models were used to correlate PC axes with latitude and longitude. To visualize continuous structure (isolation by distance) across the breeding range, we translated the average PC1 and PC2 values (based on 96 SNPs) for each sampling location to blue and green color intensity and interpolated across the breeding range using the akima R package. We then used predicted sample loadings of wintering birds (snpgdsSampLoading in SNPRelate) to assign a color for each wintering sample.

### Climate matching

For each wintering bird, we generated a probability surface to predict breeding location using the R package OriGen (Rañola *et al.* 2014). Using the 96 SNP panel, OriGen creates continuous allele frequency surfaces, which it combines to estimate the probability of an individual belonging to each grid cell on the map. We used 423 known breeding individuals to train the OriGen model (MaxGridLength=70, RhoParameter=10), then estimated the breeding location for 203 wintering birds. This produced a 70 x 23 pixel grid, with spacing approximately 1.5° longitude and 0.5° latitude, across North America with probabilities based on inferred breeding locations for each bird. We trimmed the OriGen output to the breeding range for yellow warbler and scaled the remaining probabilities to sum to 1 (Figure S2).

We obtained climate information from WorldClim (Fick & Hijmans 2017) and CRU (Harris *et al.* 2014) for every bird capture site and every grid point in the predicted breeding probability surfaces. Specifically, we averaged total annual precipitation and average annual temperature across the years 1970-2000 (WorldClim) and 1966-2015 (CRU: selected to cover the full BBS record, see below). Over the same time periods, we also obtained monthly estimates of total precipitation and monthly average, maximum, and minimum temperatures. We averaged monthly climate values across the breeding (June and July) and wintering (Nov.-Feb.) seasons (Fink *et al.* 2018). Maps of WorldClim climate values across both ranges are shown in Figure S3.

To calculate the expected breeding climate for each wintering bird (N=203), we multiplied the climate values at each grid point across the breeding region by the predicted probability of occurrence (from OriGen) at that grid point and then summed over all grid points (*i.e.,* a weighted average). To calculate climate dissimilarity between breeding and wintering sites, we first calculated the absolute value of the difference in climate values between the wintering site where the bird was captured and each site in breeding grid. Then, as before, we multiplied the climate difference values by the predicted probabilities of occurrence at each grid point and summed over all grid points. To validate our breeding climate estimates, we compared observed and predicted climate for all breeding birds, using leave-one-out cross-validation. We used linear mixed models, with sampling location as a random effect, to determine the extent to which predicted climates aligned with the true breeding site climate values.

Next, we calculated the null expectation for each bird’s breeding climate, controlling for the bird’s expected migration distance. We first calculated the geographic distance of each breeding grid point to the wintering site where each bird was captured. Using these distances and the probabilities that the bird migrated to each breeding grid cell allowed us to calculate the probability that each bird migrated different migration distances (*i.e.,* <1000km, 1000-1500 km, 1500-2000km … >6500km; Figure S2). Next, for each wintering bird, we multiplied the climate values of each grid point in the breeding range by the probability that the bird migrated that distance and then summed over all grid points. This created a null breeding climate expectation generated by migration distance probabilities (and nothing else). We used the same procedure to also calculate a null expected climate difference between the wintering and breeding sites. Finally, we computed each individual’s ‘Climate Matching Index’ by subtracting our predicted climate distance (between the wintering and breeding locations) from the null expected climate distance.

Climate Matching Index values greater than 0 indicated that an individual migrated between locations with more similar breeding and wintering climates compared to the null expectation (controlling for geographic distance). We used linear mixed models (LMMs) to determine whether the climate matching index for each climate variable was greater than 0 (*i.e.,* the intercept was significantly larger than 0). We included a random intercept of wintering location, as multiple samples often originated from the same site.

Of all the climate variables (annual total precipitation, annual average temperature, and monthly precipitation, average temperature, maximum temperature, and minimum temperature), we only found evidence that average precipitation values at breeding sites (during June and July) and wintering sites (during Nov.— Feb.) were more similar than the null expectation. To determine which birds were most likely to match monthly precipitation values between breeding and wintering sites, we added linear and quadratic effects of ‘winter site monthly precipitation’ to our model. For all models, we verified residuals were normally distributed and did not exhibit heteroskedasticity.

### Morphology

We used a published dataset (Wiedenfeld 1991) to examine morphology-climate relationships across the breeding range. This dataset includes measurements on yellow warbler museum specimens from across the entire range. We extracted only North American breeding individuals (n=145). Morphological measurements taken were: bill length, bill width, bill depth, tail length, tarsus length, sixth primary (distance from bend of wrist to the tip of sixth primary), and ninth primary (“wing length”). Although body size was not available for all specimens, tarsus length has been shown to be a reasonable proxy (Senar & Pascual 1997). We therefore scaled all other measurements by tarsus length by calculating residuals from simple linear regressions. We used a generalized least squared model to examine correlations between each of these morphological measurements and precipitation with an exponential correlation structure accounting for longitude and latitude (*i.e.,* spatial autocorrelation). P-values were adjusted for multiple tests using a false discovery rate correction.

### Demography

To examine temporal relationships between precipitation and abundance, we used data from the North American Breeding Bird Surveys (BBS) (Sauer *et al.* 2017). First, we used hierarchical models to estimate the effect of temporal fluctuations in precipitation on yellow warbler abundances. We divided the breeding range into 100km hexagonal grid cells (n = 1240) and identified the grid cell for each BBS route where at least one individual had been observed (n = 3710). For each BBS route, we then calculated the deviation from the mean precipitation (from 1968 to 2015) for each year. We focused on precipitation deviation in the year prior to the BBS survey, as surveys are done in the spring/summer and same-year precipitation would thus include conditions not yet experienced. Because there can be lag effects of climate on demography, we also analyzed precipitation from two years prior. We used hierarchical models to estimate effects of precipitation (1 and 2 year lags) on yearly variation in abundance as described in (Link & Sauer 2002), with the exception of using a negative binomial to model count data and including the 1240 hexagonal grid cells as the ‘strata.’ Models were run using JAGS in the R environment, with 4-chains each run for 50K iterations, with 20K discarded as burn-in, with a thin-rate of 120, yielding a total of 1000 posterior samples. We assessed convergence by visually inspecting traceplots of the chains, and verifying that the Gelman-Rubin Rhat statistic (Gelman & Rubin 1992) was less than 1.1 for all parameters.

Finally, we explicitly tested the hypotheses that the relationship between abundance and precipitation varied over geographic space by using a spatial spline GAM model. Some grid cells’ estimates were more precise while others had substantial uncertainty in their posterior distributions, so we propagated uncertainty in the estimates of the climate-population relationship to ensure that false precision was not influencing our result. To do so we weighted each grid cells’ posterior mean estimate of the effect of precipitation by the inverse of the squared standard deviation of the posterior (equivalent to the inverse variance, or precision of the estimate). Finally, we repeated the analysis with grid cell sizes of 200km and 400km to ensure our results were robust to the choice of grid cell size.

## Results

Genetic variation across the yellow warbler breeding range showed strong patterns of isolation by distance. Principal components analysis of 104,711 SNPs derived from RAD-Seq data for 195 breeding individuals (Fig S4) showed clear associations with longitude and latitude (Figure S4; PC1 vs longitude: *R*^*2*^=0.86,*p*<0.001; PC3 vs. latitude: *R*^*2*^=0.44,*p*<0.001). RAD-Seq data were used to identify a set of 192 SNPs, of which 157 yielded high-quality genotypes in an additional 224 breeding birds to produce the spatial map of population structure we used to visualize migratory connectivity. Although the Evanno method identified K=2 as the optimal number of populations, we still see obvious geographic patterns at higher values of K, likely driven by strong isolation by distance (Figures S5-S9) (Bradburd *et al.* 2018). The highest K value at which each group has individuals with majority ancestry (>50%) was K=5 (Figure S3). Of the 419 breeding birds remaining after quality filtering, 83% (349) had at least 50% ancestry from a single K=5 group.

Although K=2 may more correctly identify barriers to gene flow, the K=5 scenario provides higher resolution for visualizing migratory connectivity while still accurately assigning individuals to breeding groups (Figure 1). Cluster-based assignment has similarly been successfully used to describe movement patterns in fisheries systems with low population structure (Layton *et al.* 2020; Spies *et al.* 2020). Assignment to the five groups, based on a set of 96 SNPs genotyped using Fluidigm SNPtype assays, was robust: 87.6% of breeding samples were correctly assigned. We were able to assign 189 wintering individuals to the five breeding groups. We found that birds from the four westernmost groups all winter broadly across Central America. Nevertheless, there was substructure in the wintering range within Central America; for example, the wintering range of individuals that breed in the southwestern US extends further north within Mexico than do the other groups. Individuals breeding in central North America were found throughout Central America, but also in South America. Birds from the Eastern US were only found wintering in South America (not in Central America). Importantly, some of these connections were obscured when describing connectivity with only two groups (K=2). Namely, the strong connection between the eastern population and South America is not clear at that resolution (Figure S10). Using PCA to map migratory connectivity in a more continuous matter resulted in a qualitatively similar picture (Figure S11).

**Figure 1.**
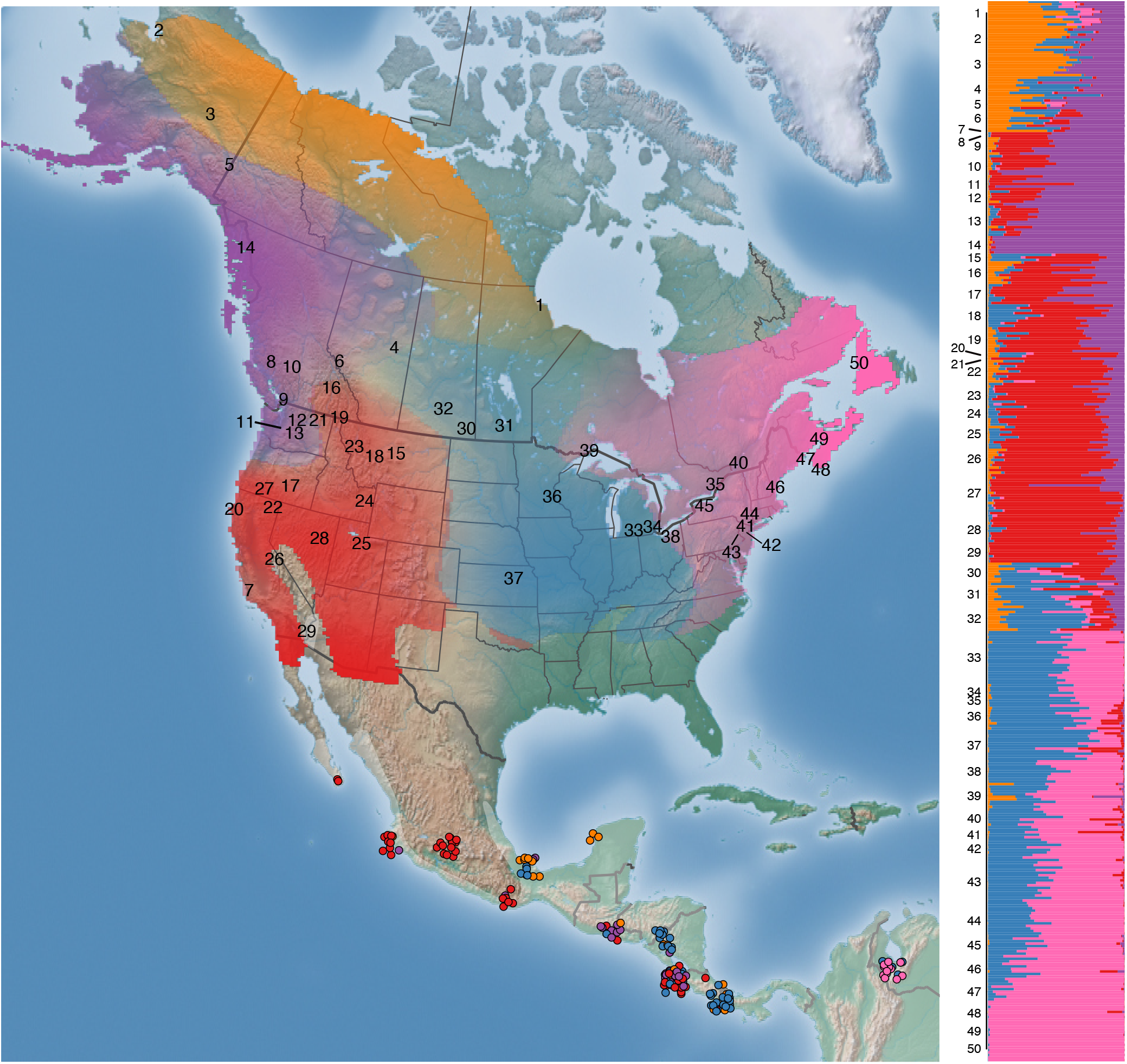
Migratory connectivity in yellow warbler. North American breeding range shows the five groups inferred by STRUCTURE (K=5), spatially interpolated across the range (ancestry for the dominant group in a particular location is represented by transparency). The barplot on the right shows the structure ancestry plot with numbers corresponding to the same numbers on the breeding range of the map. Points on wintering grounds are colored based on breeding population assignment.

To test whether individuals track climate between breeding and wintering grounds, we also estimated specific predicted breeding locations (*i.e.,* latitude and longitude) for each wintering individual. In general, there was agreement between the predicted location and the predicted group based on STRUCTURE analysis (Figure S12). From cross validation with breeding birds, we found that predicted and observed climate values were strongly correlated (conditional *R*^*2*^ 0.59-0.9), suggesting high accuracy in our predicted breeding climate (Figure S13). We calculated a “Climate Matching Index” (CMI), defined as the degree to which wintering and inferred breeding climate were more similar (CMI>0) or less similar (CMI<0) than expected given a null model accounting for migratory distance. We found that birds migrated between areas with similar precipitation regimes, specifically areas with similar amounts of monthly precipitation during the times of year when the species is present (Table S3; Figure 2A,C). This pattern was strongest in birds wintering in the driest regions of Central America and breeding in the driest regions of North America (Table S4; Figure 2B). These results were robust to the choice of climate dataset (CRU vs. Worldclim; Table S3, S4; Figure S14) and randomizations show that our results are robust to the levels of uncertainty in estimates of breeding climates (Figure S15). We did not find any evidence of individual-level tracking for mean, maximum, or minimum temperature or when temperature and precipitation were calculated over the entire year (*i.e.,* including periods when the species is absent; Table S3; Figure S16).

**Figure 2.**
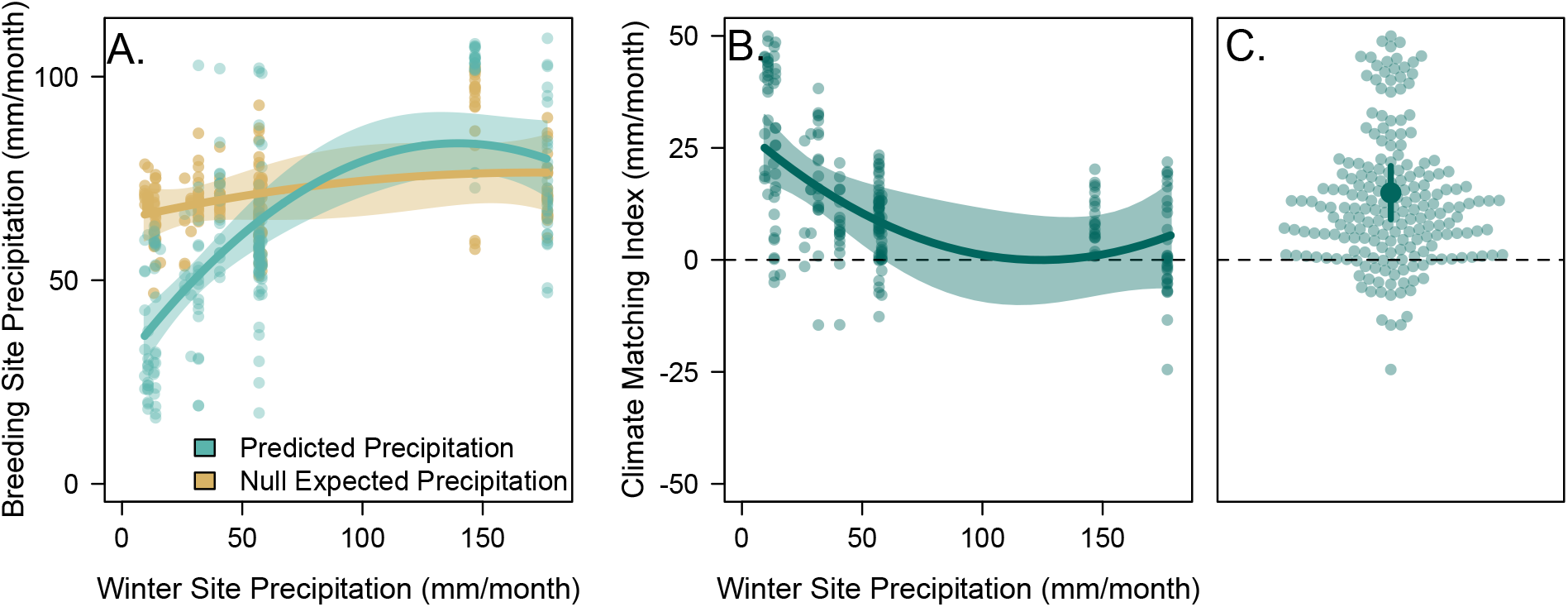
Individual climate tracking in yellow warblers caught in the wintering range. (A) Individuals captured at drier wintering sites were predicted to breed at drier sites (based on genetic analyses; blue points and lines). This trend that did not occur under the null expectation, in which breeding locations were only constrained by migration distances (brown points and lines). (B) After accounting for migration distances, the Climate Matching Index (see methods) indicates similarity between breeding and wintering climate is highest for birds in dry regions. (C)The Climate Matching Index was significantly greater than 0, providing evidence for individual-level climate tracking of precipitation regimes. Precipitation values were calculated from CRU data. In (A) and (B), lines represent predictions from mixed-effects models, shaded regions are 95% confidence intervals, points are individual birds. In (C), transparent points are individual birds, the solid point is the mean estimated Climate Matching Index, and the line is the 95% confidence interval.

Adaptation across climate gradients could drive divergence in morphological traits, as the link between climate and morphology has been demonstrated in a number of avian systems (Grant & Grant 1993; Chavarria Pizarro *et al.* 2019). We found that, in specimens collected across the breeding range (n=145), bill length (*F*=14.04, adjusted p=0.001) and bill depth (*F*=8.29, adjusted p=0.025) were positively correlated with breeding season precipitation (Figure 3). Specifically, birds with longer, deeper bills were found breeding in wetter regions. No other morphological traits were correlated with breeding precipitation (Figure S17).

**Figure 3.**
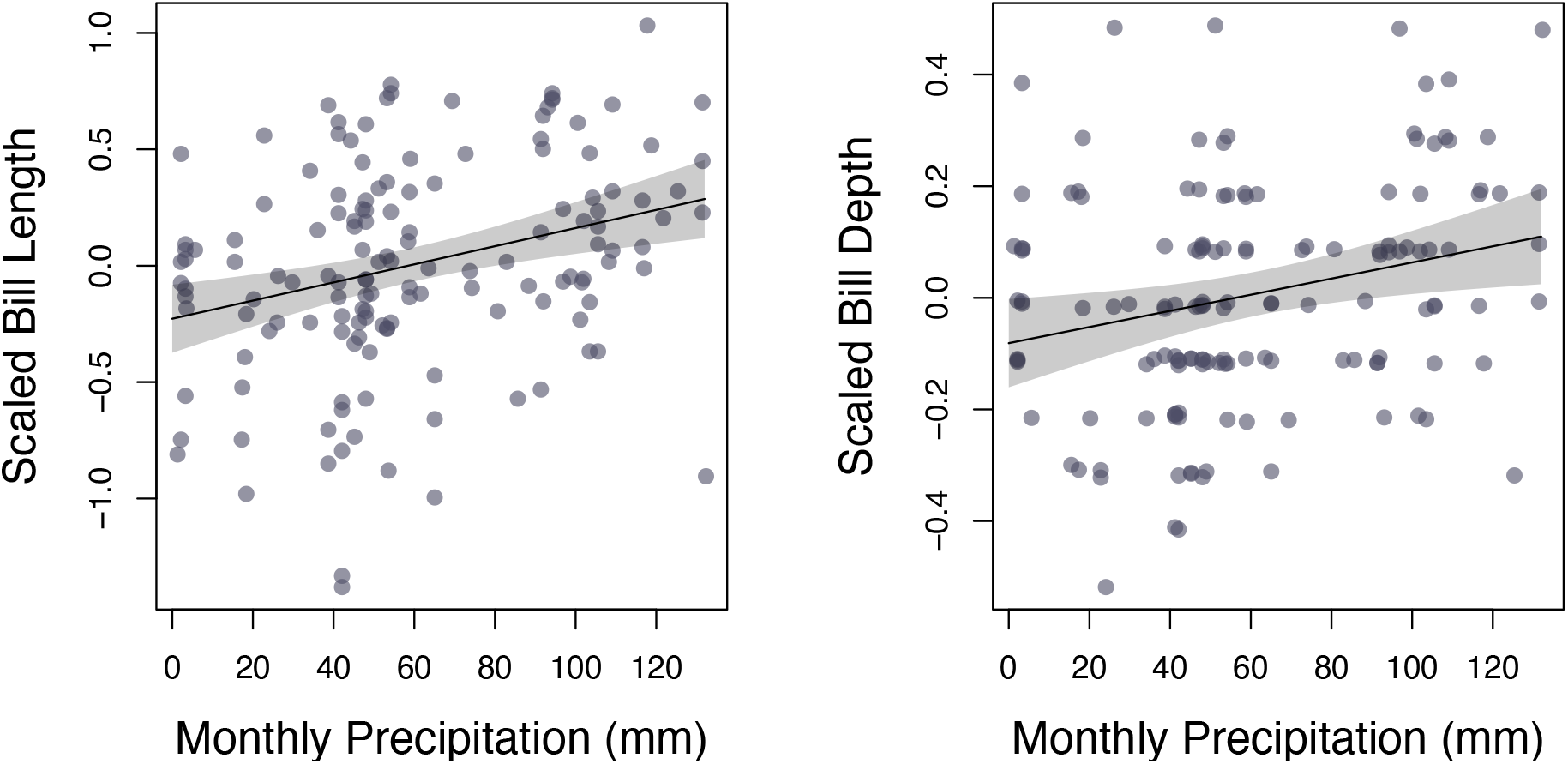
Correlation between bill length (A) and bill depth (B) and precipitation for birds sampled across the breeding range. Measurements were scaled by tarsus length as a proxy for body size. Morphological data are from Weidenfeld (1991). Precipitation values were calculated as average precipitation during the breeding season, extracted from the CRU dataset.

**Figure 4.**
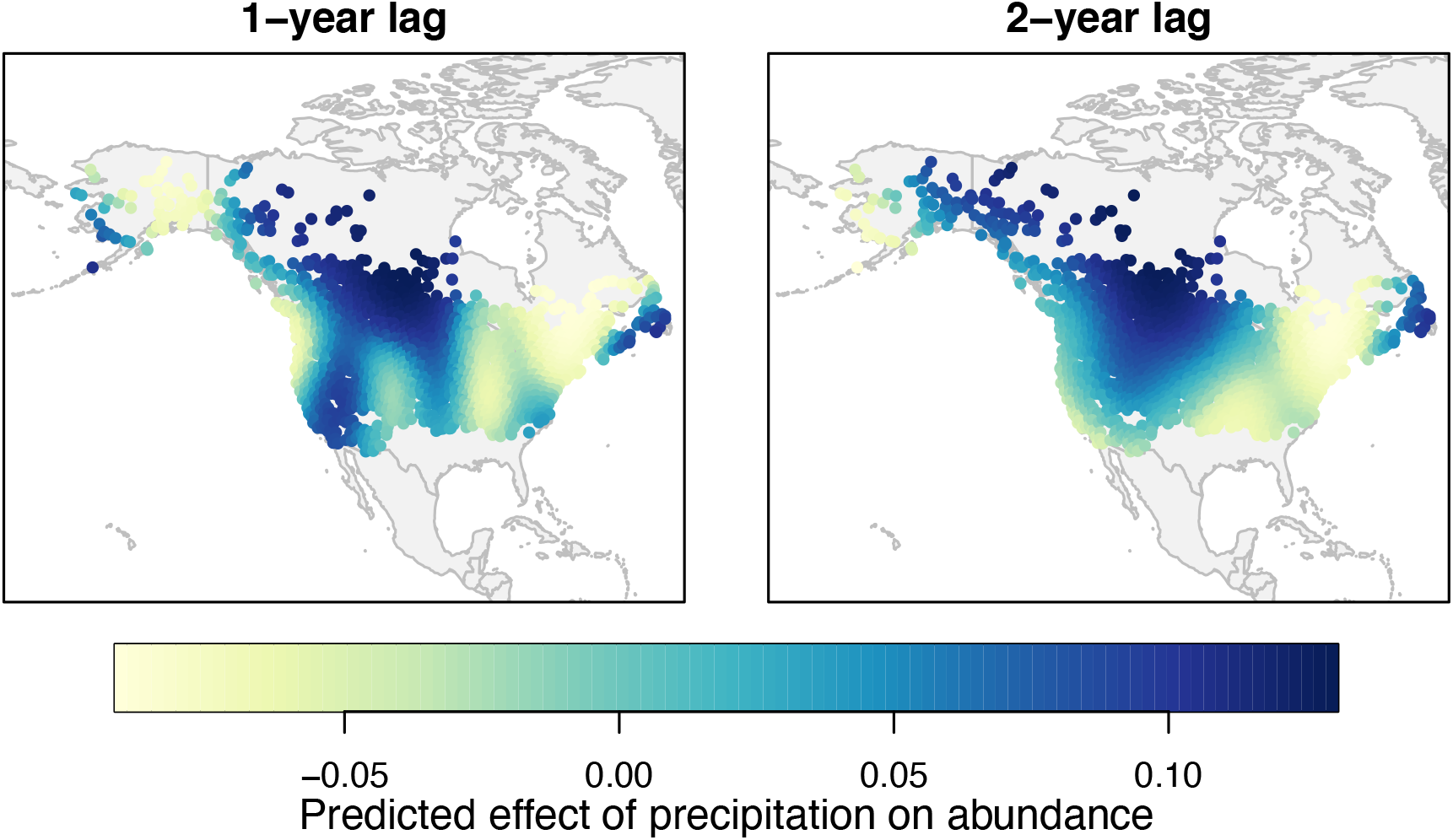
Associations between precipitation anomaly and yellow warbler relative abundance across the breeding range. Points represent 100km grid cells across sites where the species was detected in Breeding Bird Surveys. Colors indicate the predicted effect of precipitation anomaly on abundance (*i.e., β* values from a Bayesian hierarchical model) from a GAM (see methods). Left plot shows associations with climate for the calendar year prior to the survey and right plot shows two years prior.

Temporal fluctuations in population size associated with climate could provide opportunities for selection of climate-adapted genotypes if such events result in differential survival and/or reproduction. Using abundance data from the North American Breeding Bird Survey (BBS), we found significant spatial variation in the relationship between abundance and precipitation in the year prior to the survey (spatial spline term p<0.001) as well as two years prior (p<0.001) Generally, the positive effects of precipitation on abundance were in western areas (which are often drier overall), while negative effects were largely seen in the wetter eastern regions. Qualitative patterns of the relationship over space (Figure S18), and significance of the spatial spline term (all p<0.05), were robust to alternative grid size choices.

## Discussion

Predicting species response to climate change will require an understanding of the extent to which climate constrains the ranges of genotypes, individuals, and populations (Fitzpatrick & Keller 2015; Bay *et al.* 2018). For migratory species, this is especially challenging. Overlap between climate niches on breeding and wintering grounds at the species level can be examined using observational data, but similarity among breeding and wintering climates at the individual level requires the ability to pinpoint breeding and wintering locations accurately for many individuals from across the range. We capitalize on innovations in genetic sequencing and analysis to accomplish this level of resolution, showing that yellow warbler individuals track similar precipitation (but not temperature) regimes between the breeding and wintering seasons. Coupled with associations between precipitation and both morphological (Figure 3) and genetic (Bay *et al.* 2018) variation, our results provide a potential mechanism for the maintenance of local adaptation across the annual cycle.

A prerequisite for continued, reinforced natural selection across the annual cycle is site fidelity. If individuals move to new breeding or non-breeding locations each year, signals of selection might be lost. Our study and others found strong isolation by distance in the breeding range (Gibbs *et al.* 2000), suggesting that individuals migrate to the same regions year after year. Multi-annual bird banding studies support this observation (Cilimburg *et al.* 2002). In the wintering range, we found mixing of our five groups, but also some sorting, with western populations wintering farther north in Mexico and eastern populations only found in South America. These findings support previous microsatellite and stable isotope studies of migratory connectivity in yellow warbler, which have reported segregation between eastern and western lineages on their wintering grounds (Milot *et al.* 2000; Boulet *et al.* 2006). The higher resolution provided by RAD-Seq data allows us to further refine the subtle differences in wintering grounds among breeding groups. Unlike a recent geolocator study (Witynski & Bonter 2018), we find little evidence for crosswise migration, although such patterns could persist within our breeding groups. Although migratory connectivity may not appear extremely strong when examined at the population level, that does not preclude the possibility that individual birds within each of the five groups reliably use the same breeding and wintering grounds. While we find that climate explains migratory connectivity beyond what is expected by distance alone, we cannot exclude potential confounding forces shaping the evolution of migratory routes. Additionally, it is important to note that the spatial scale of climate data coupled with the uncertainty in predicting breeding location does not allow us to examine variation associated with microclimate, but rather associations with broad regional climate regimes. Other selective mechanisms might drive genetic and morphological divergence at smaller scales. The combination of genetic and morphological associations with precipitation across the breeding range, as well as climate tracking across the annual cycle suggests that adaptation across climate gradients might also exist on the wintering grounds. However, data on the distribution of adaptive genetic and phenotypic variation and the relationship of such traits to climate during the wintering period is still needed.

Local climate can impact bird populations through multiple pathways; for example, by directly affecting physiology or through indirect effects on resources. We find a correlation between bill size (both length and depth) and precipitation. Interestingly, in the resident subspecies of yellow warbler found in Costa Rica, the mangrove warbler (*S. petechia xanthotera*), bill size also increases with precipitation (Chavarria Pizarro *et al.* 2019). There are several potential mechanisms through which different precipitation regimes could select for different bill sizes. One possibility is that larger bills more efficiently dissipate heat. Although this has largely been explored in connection with temperature alone, there is some evidence that larger bill sizes may be adapted to more humid environments where evapotranspiration is less efficient (Gardner *et al.* 2016) and that precipitation and temperature interact to shape selection on bill size (LaBarbera *et al.* 2020). Another hypothesis is that variation in bill size is a response to food availability, a pattern which has been observed in a wide variety of avian taxa (Grant & Grant 1993; Badyaev *et al.* 2008; Bosse *et al.* 2017). As precipitation can structure the size distribution of insect communities (Janzen & Schoener 1968), this might lead to different optimal bill morphologies. Although our study suggests that bill size may be under selection across precipitation gradients in the breeding range, alternative explanations, including neutral processes or phenotypic plasticity cannot be ruled out at this time.

Selective drivers can affect demographic trends through spatial and temporal variation in mortality and fecundity. For example, Sillet et al. (Sillett 2000) found lower fecundity of black-throated blue warblers (*Setophaga caerulescens*) in El Niño years in the eastern US, with fecundity correlated with prey biomass. Similarly, yellow warbler populations in Manitoba, Canada had lower survival and reproduction during El Niño years (Mazerolle *et al.* 2005). The opposite pattern was observed in a study across 10 landbird species in the Pacific Northwest, with higher reproductive success in El Niño years (Nott *et al.* 2002). Indeed, models suggest that El Niño years are associated with higher survival in this region (LaManna *et al.* 2012). These studies highlight not only the effects of climate on survival and reproduction in migratory birds, but also the potential geographical variation in demographic effects of climate anomalies. Correspondingly, we found that the relationship between precipitation anomaly and abundance was sometimes strong but varied across the breeding range. In parts of the western US, which experiences the driest breeding climates, wetter years were followed by higher yellow warbler abundance. In contrast, we saw negative effects of precipitation in some eastern regions. As fluctuations in abundance are a function of survival and fecundity, they represent an opportunity for selection. Differential fitness following the driest years, when abundances are lowest in the western population, could lead to increased frequency of particular phenotypes favored in dry climates, shaping the genotype-phenotype-environment variation we observe across the breeding range. Of course, many non-climatic factors across both the breeding and wintering grounds also affect populations trends. Further work on the mechanistic link between phenotype, fitness, and climate will help elucidate the extent to which fluctuations in abundance result in selective shifts.

Our results suggest that individuals sort non-randomly across breeding and wintering ranges so as to minimize differences in precipitation regimes. On the species level, this “niche tracking” has been suggested as an explanation for the evolution of migration – as an escape from seasonality (Winger *et al.* 2019). It is not clear how a similar explanation would play out at an individual or population level. We propose that for species that have already established migratory life histories, it may be beneficial that selection on traits that are adaptive in one part of the life cycle be reinforced across all seasons. One might imagine that birds with traits that allow them to persist in the driest breeding conditions then become the most fit in the driest wintering regions if the mechanisms of selection are either the same or parallel across both regions. Coupled with strong migratory connectivity, this would lead to intraspecific correlations between climate on the breeding and wintering grounds, as we observe here.

In conclusion, our study provides evidence that precipitation regimes (but not temperature) structure genetics and morphology, migratory connections, and population trajectories in a widespread migratory bird species. Such information about the ecological and evolutionary responses to changes in climate regime will be essential for understanding and predicting response to future global change.

## Acknowledgements

We thank many people who assisted in sample collection, especially M. Boulet, R. Dawson, E. Milot, K. Hobson, H. L. Gibbs, B. Keith, the University of Washington Burke Museum, S. Albert, T. Kita, and the many Institute for Bird Populations and MAPS volunteers for providing or assisting with collection of samples. This work used the Extreme Science and Engineering Discovery Environment (XSEDE), which is supported by National Science Foundation grant ACI-1548562. This work was made possible by an NSF Postdoctoral Fellowship to R.A.B, and support from NSF Rules of Life EAGER to R.A.B. (1837940), National Geographic to K.R. (WW-202R-17), California Energy Commission to K.R. (EPC-15-043), NSF CAREER to K.R. (1942313) and First Solar Incorporated. Finally, we thank three anonymous reviewers for comments that substantially improved the manuscript.

## Notes

### Competing Interest Statement

The authors have declared no competing interest.

