## Supplementary material for "Genetic variation reveals individual-level climate tracking across the annual cycle of a migratory bird": Figure S1, Figure S2, Figure S3, Figure S4

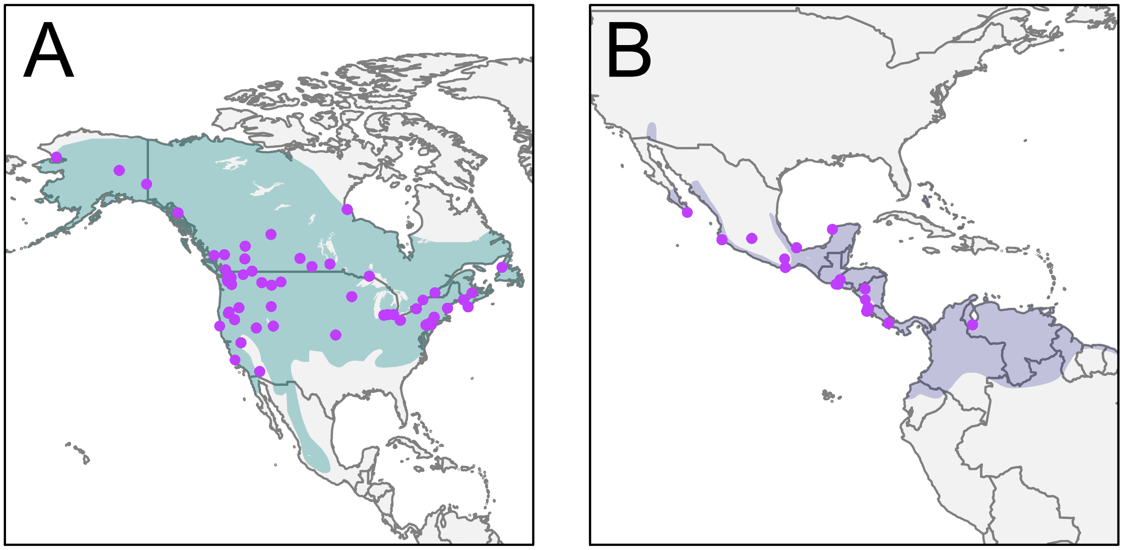


**Figure S1.** Samples used for population structure and connectivity mapping. Maps show samples from breeding (A), and wintering (B) birds. Breeding and wintering ranges are shown in green and purple, respectively. All samples used for analysis are listed by sampling location in Table S1.

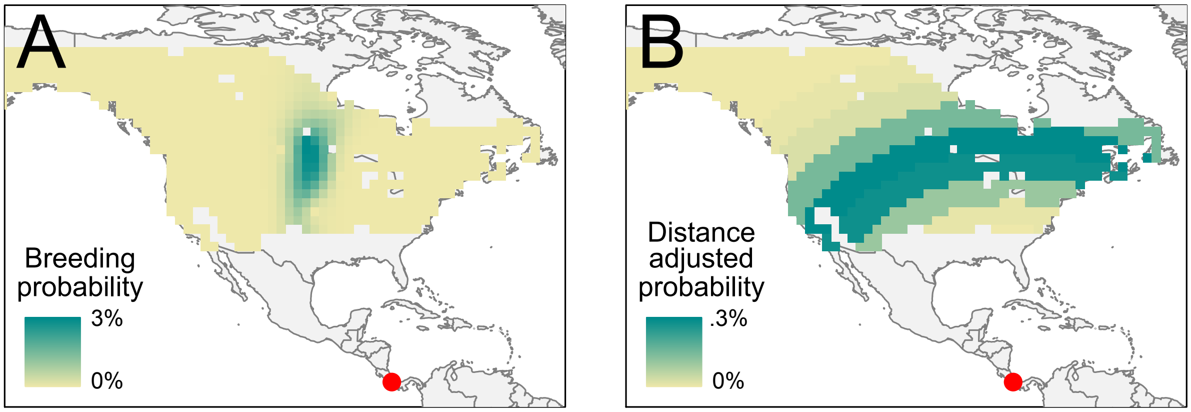


**Figure S2.** Example breeding probability surface and null surface used for calculating the climate matching index. Here we show data from a single individual sampled in Costa Rica during the wintering season. (A) shows the probability of occurrence during the breeding season for each pixel across the breeding range calculated from genetic data while (B) shows the distance-corrected null probability. These two surfaces are used to estimate the expected and null climate distances, the difference between which is the climate matching index (CMI).

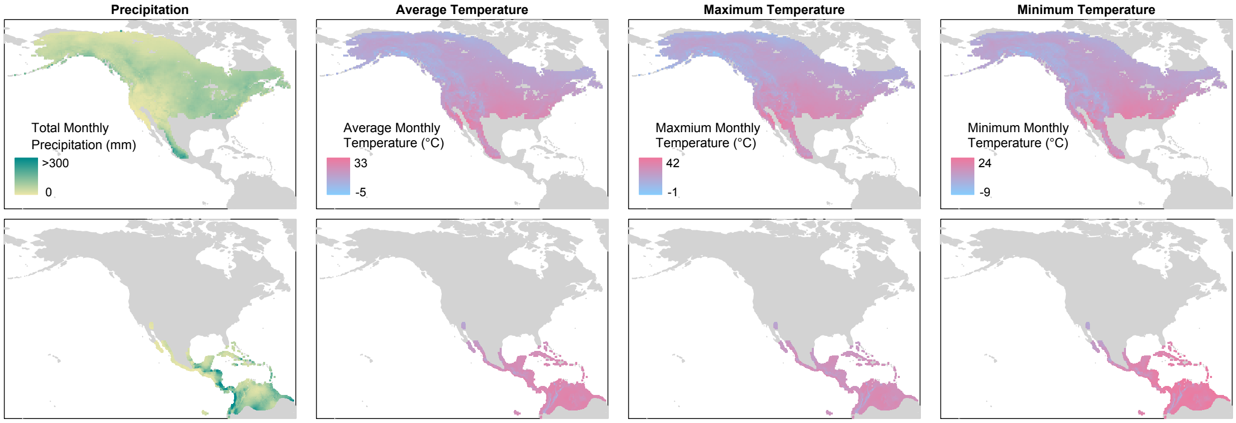


**Figure S3.** Climactic variation across the breeding and wintering ranges of yellow warblers. Here we show precipitation as well as average, maximum, and minimum monthly temperature during the relevant months: June-July for the breeding range and November-February for the wintering range. Climate data was downloaded from the WorldClim database.

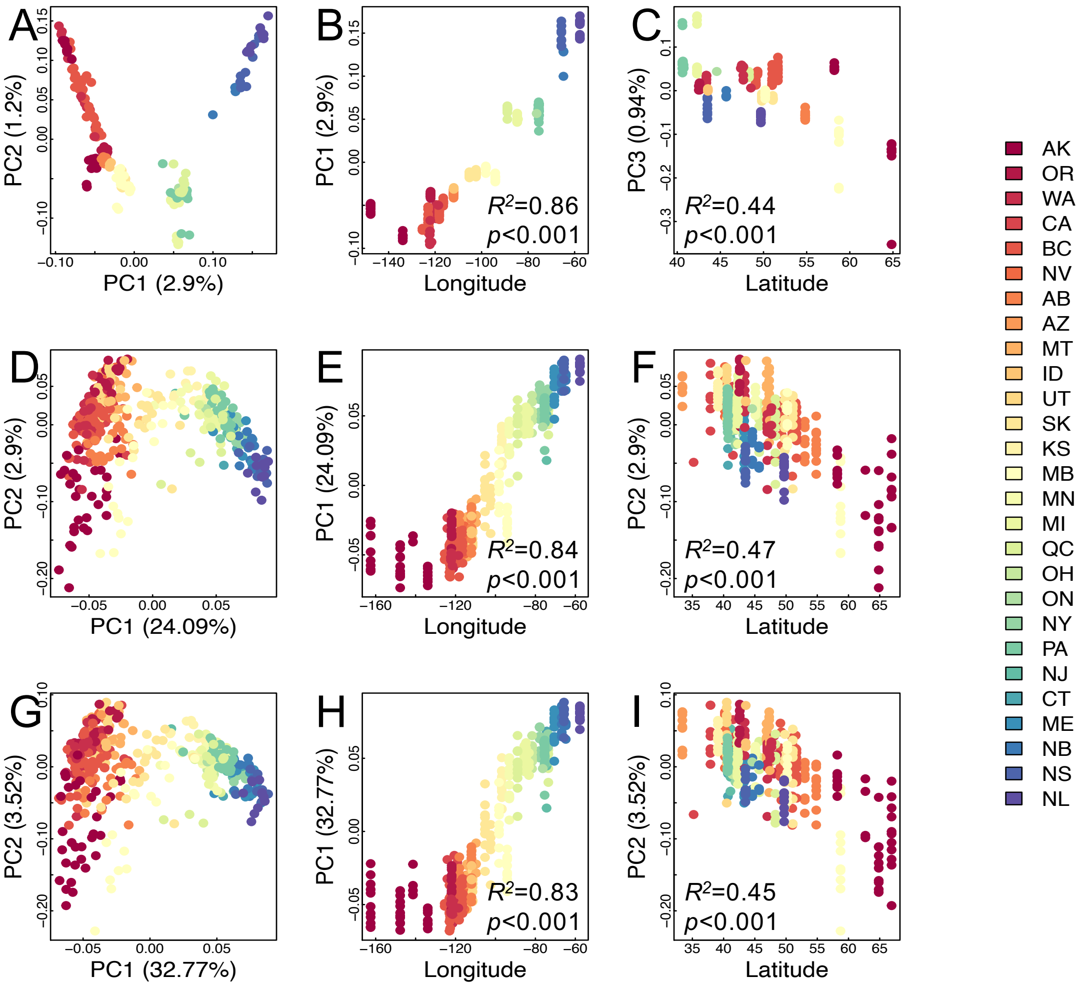


**Figure S4.** Principal components analysis using all 104,711 SNPs (A,B,C) and using 157 (D,E,F) or 96 (G,H,I) SNPs chosen for Fluidigm SNPtype assay. Plots show first PC biplot of first two PC axes (A,D,G) as well as associations between PC axes, longitude (B,E,H) and latitude (C,F,I). Samples are colored according to breeding state and states are ordered based on mean longitude.

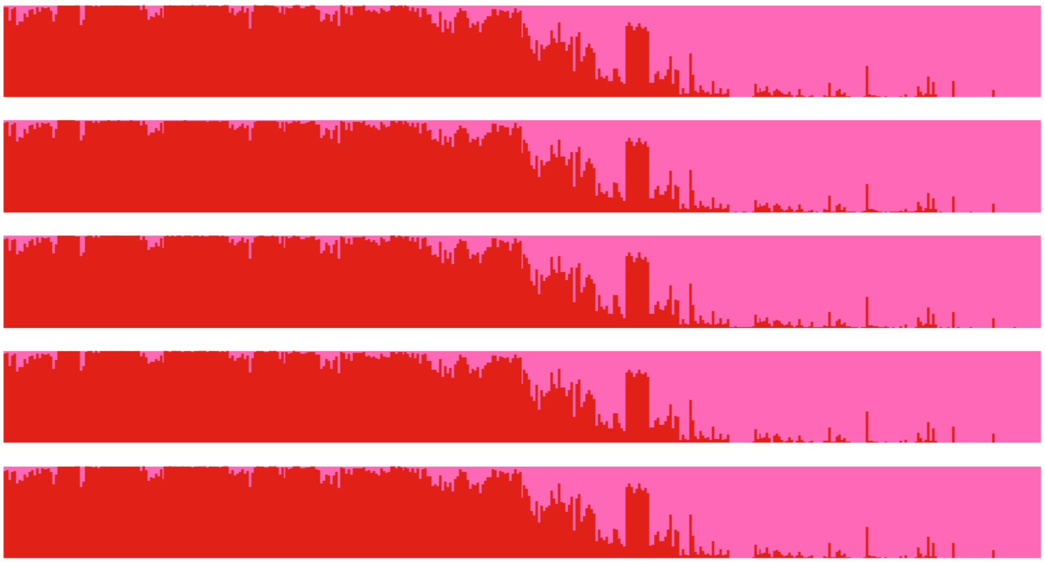

**Figure S5.** Five replicate runs (K=2) of STRUCTURE across 419 breeding yellow warbler samples genotyped at 157 SNPs. Individuals are ordered by longitude.

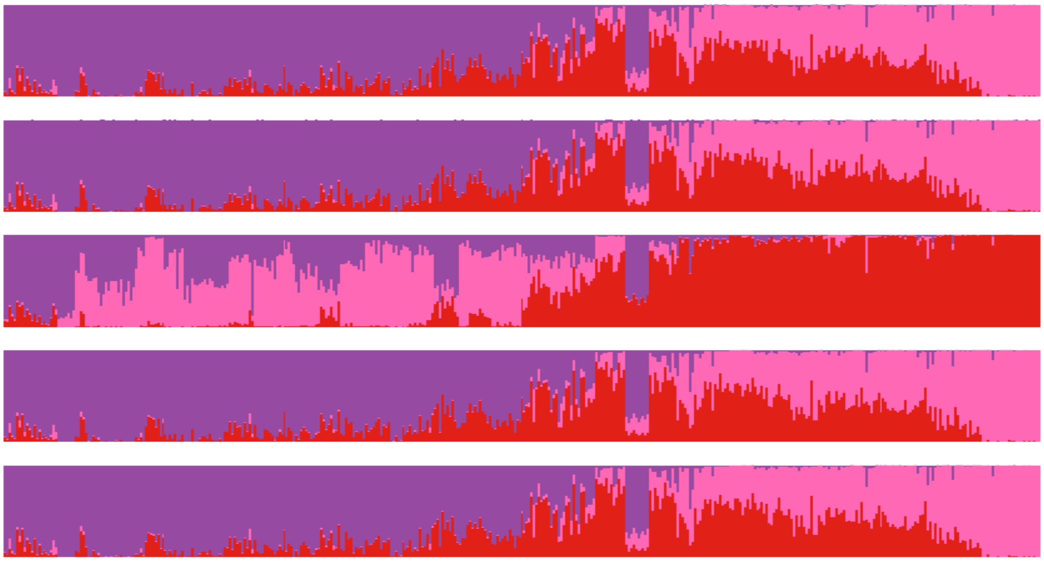

**Figure S6.** Five replicate runs (K=3) of STRUCTURE across 419 breeding yellow warbler samples genotyped at 157 SNPs. Individuals are ordered by longitude.

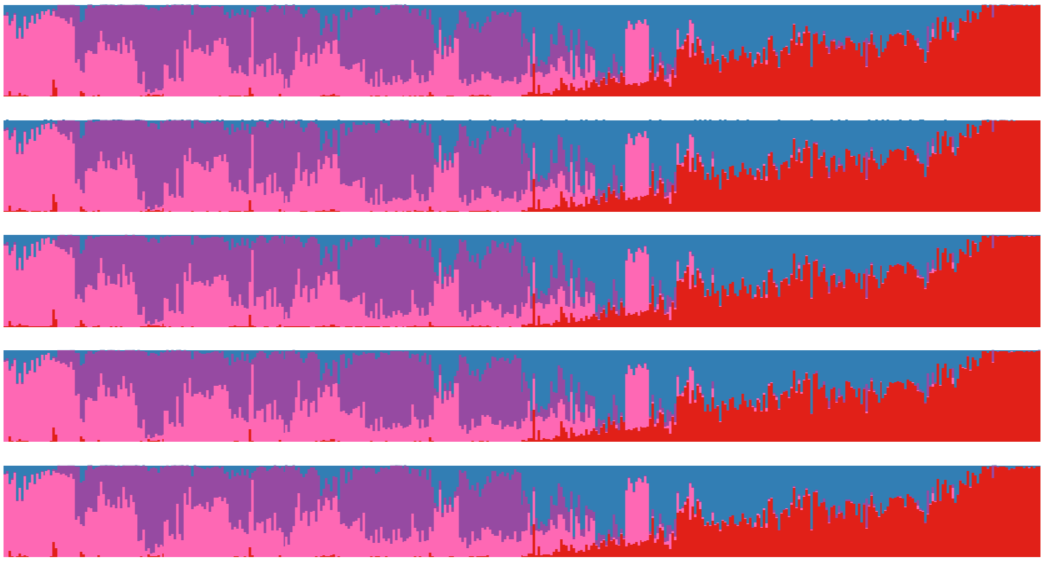

**Figure S7.** Five replicate runs (K=4) of STRUCTURE across 419 breeding yellow warbler samples genotyped at 157 SNPs. Individuals are ordered by longitude.

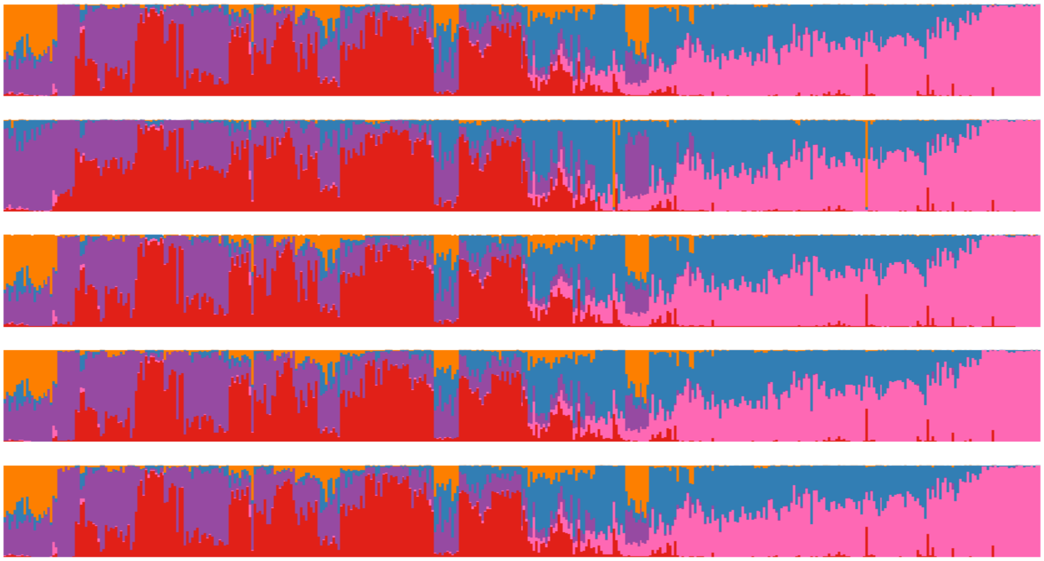

**Figure S8.** Five replicate runs (K=5) of STRUCTURE across 419 breeding yellow warbler samples genotyped at 157 SNPs. Individuals are ordered by longitude.

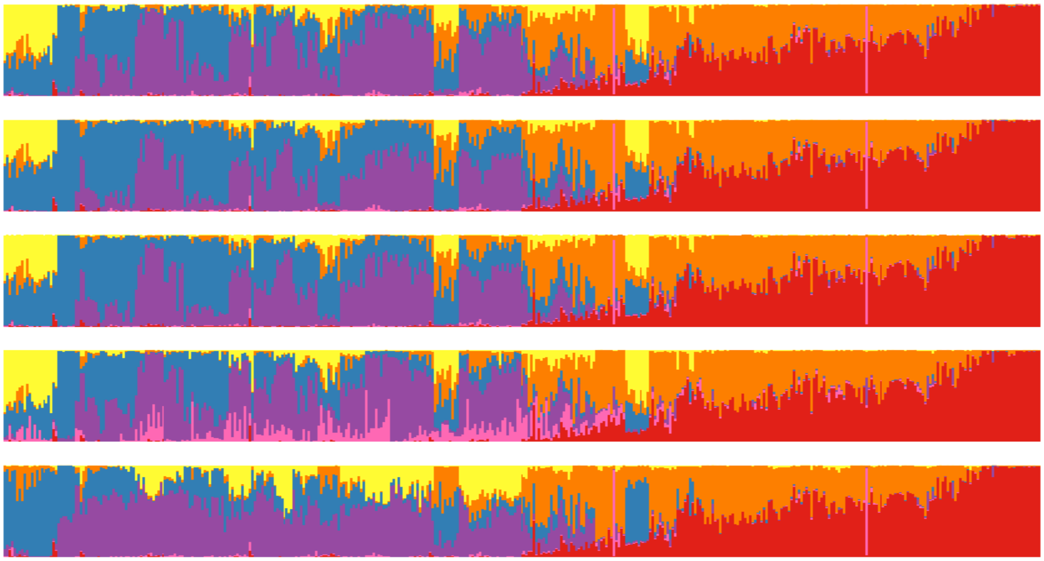

**Figure S9.** Five replicate runs (K=6) of STRUCTURE across 419 breeding yellow warbler samples genotyped at 157 SNPs. Individuals are ordered by longitude.

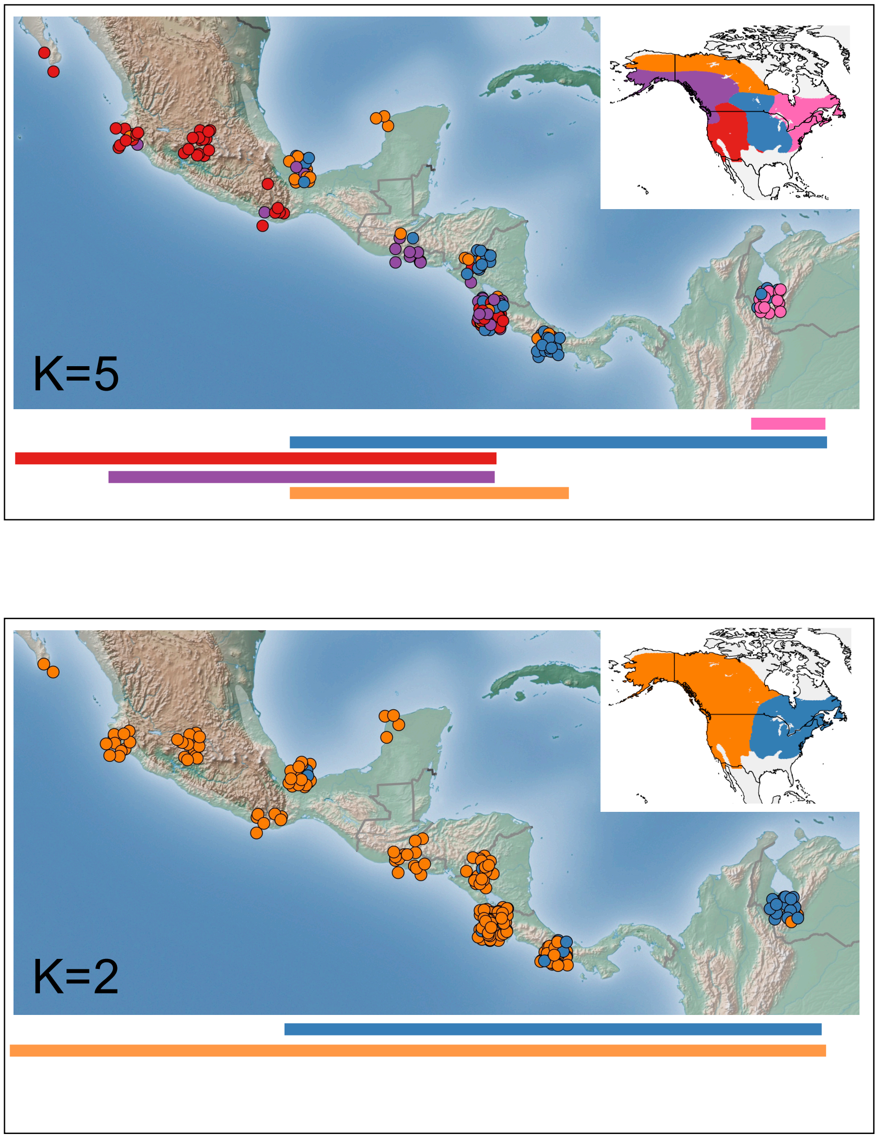


**Figure S10.** Comparison of migratory connectivity based on STRUCTURE runs with K=2 vs. K=5. Points are based on rubias assignments to breeding groups (colors shown in inset map). Lines below the map show the longitudinal ranges on the wintering ground for each of the breeding groups.

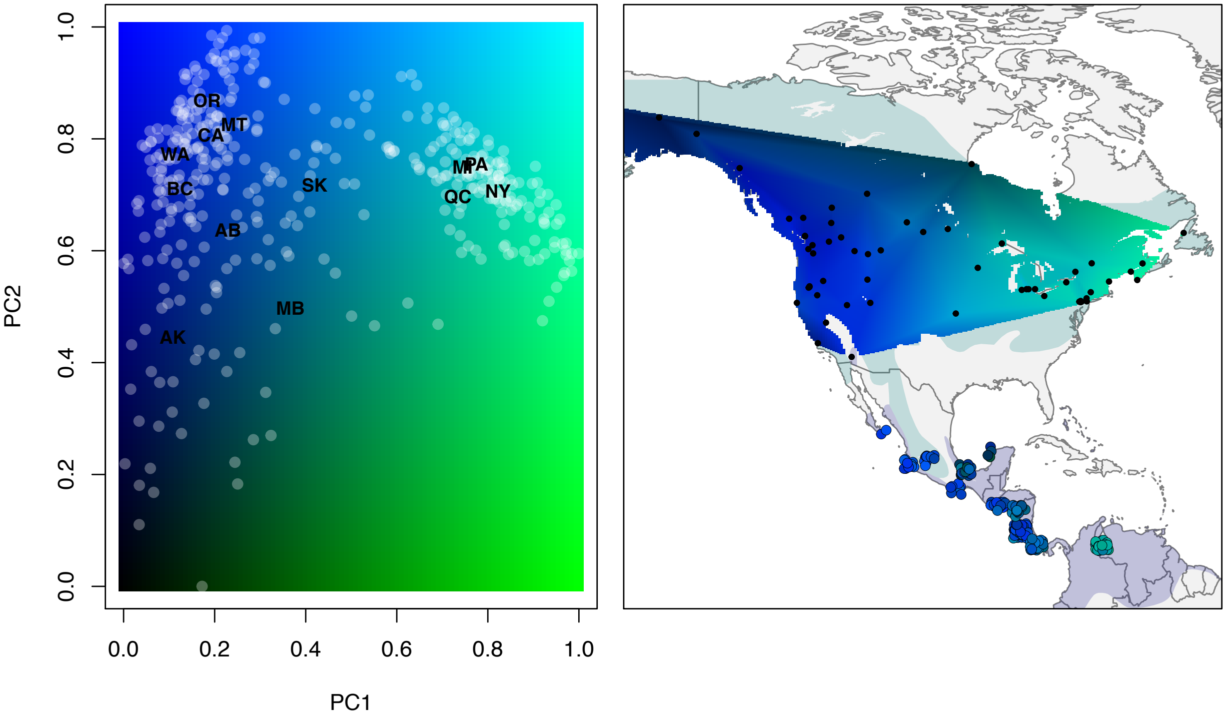


**Figure S11.** Migratory connectivity based on a continuous surface defined by principal components loadings. Left plot shows the locations of individuals and select states on the PCA with PCA loadings defining blue (PC2) and green (PC1) color intensity. Color intensities are spatially interpolated across the breeding range in the figure on the right. Wintering birds are assigned a color based on their predicted loadings on PC1 and PC2.

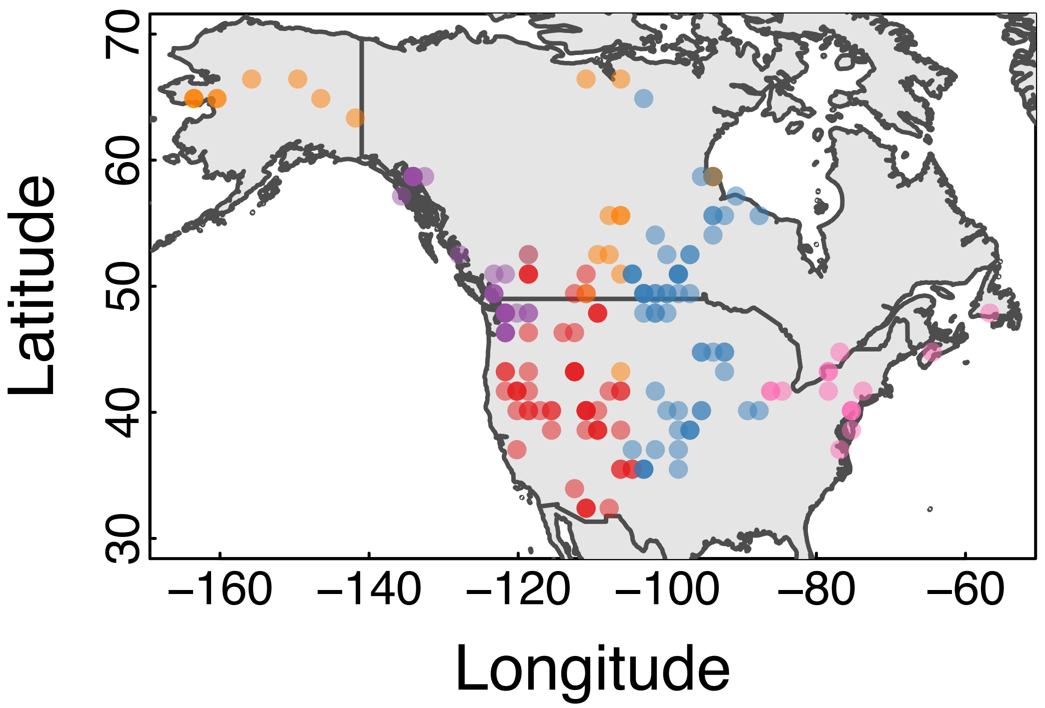


**Figure S12.** Concordance between breeding range group assignments (rubias) and continuous location assignments (OriGen). Each point represents a bird caught on the wintering range. The color of the point represents the group assignment and the location of the point shows the pixel with the highest probability of occurrence for that individual.

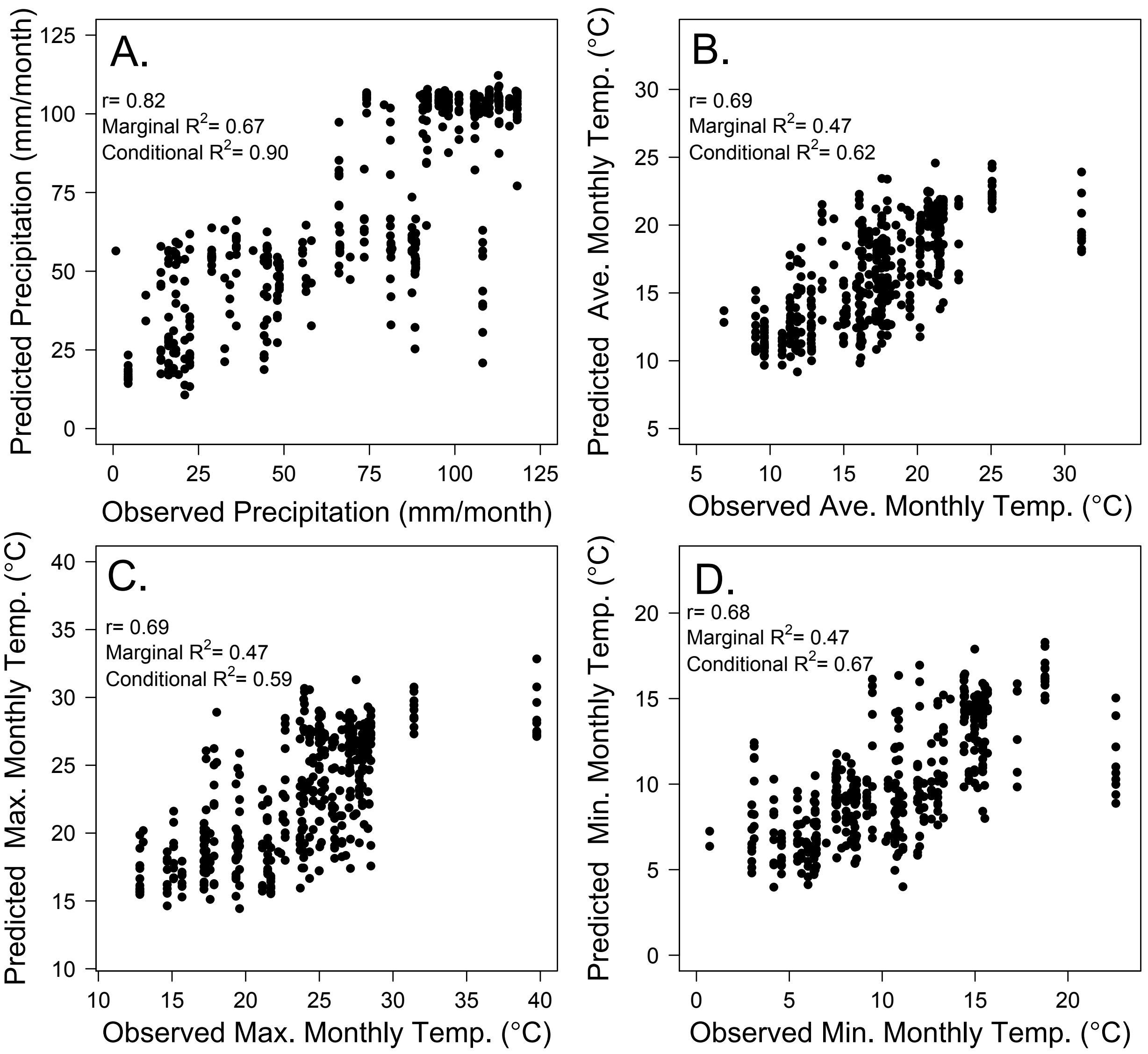


**Figure S13.** Accuracy of climate predictions based on OriGen probability surfaces. Here, we show for all birds caught in the breeding range the climate value for the site at which the bird was caught (x-axis) compared to the expected breeding climate calculated by multiplying the probability of occupancy for each pixel by the climate for each pixel. Using data from the CRU database, we show strong correlations between observed and predicted climate for monthly precipitation (A), average monthly temperature (B), maximum monthly temperature (C), and monthly minimum temperature (D) during the breeding months.

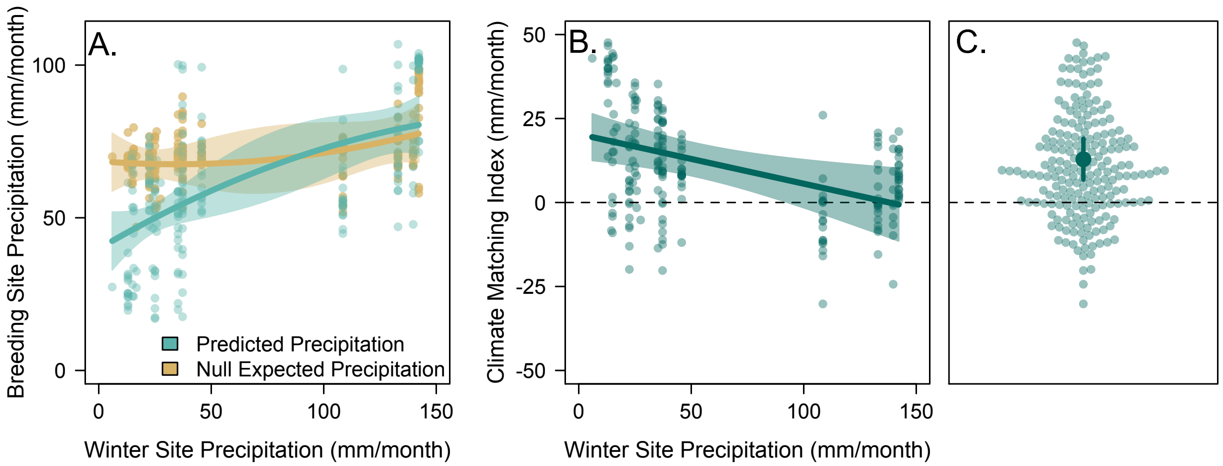


**Figure S14.** Individual climate tracking in yellow warblers caught in the wintering range, using precipitation data from Worldclim*.* (A) Individuals captured at drier wintering sites were predicted to breed at drier sites (based on genetic analyses; blue points and lines). This trend did not occur under the null expectation, in which breeding locations were only constrained by migration distances (brown points and lines). (B) After accounting for migration distances, the Climate Matching Index indicates similarity between breeding and wintering climate is highest for birds in dry regions. (C)The Climate Matching Index was significantly greater than 0, providing evidence for individual level climate tracking of precipitation regimes. In (A) and (B), lines represent predictions from mixed effects models, shaded regions are 95% confidence intervals, points are individual birds. In (C), transparent points are individual birds, the solid point is the mean estimated Climate Matching Index, and the line is the 95% confidence interval.

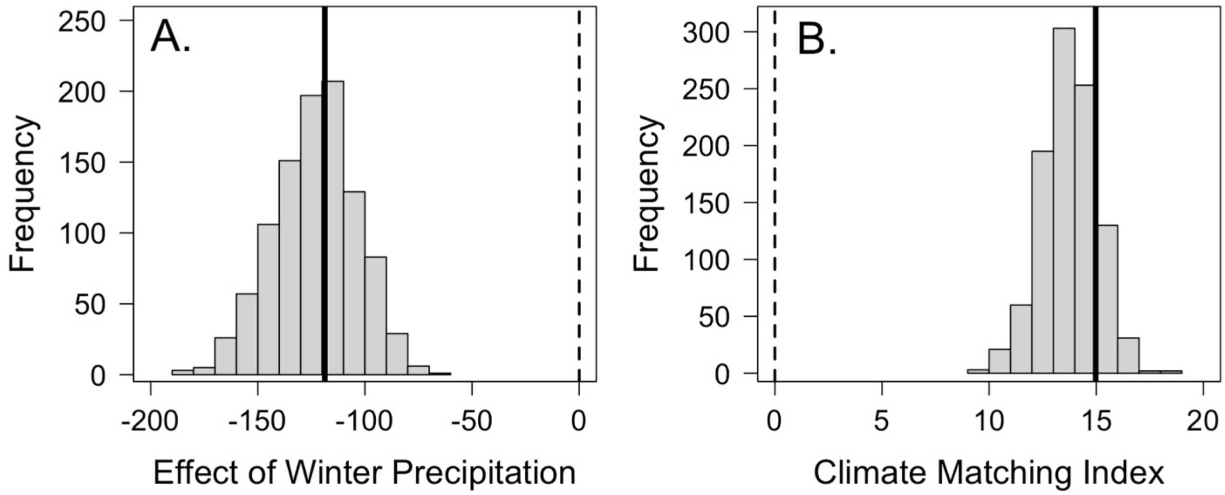


**Figure S15.** Randomizations show that climate matching inference is robust to uncertainty in prediction of breeding climate. For each wintering bird, we drew a breeding site based on the spatial probability distribution from OriGen and calculated climate matching indices based on these values. We conducted 1000 bootstraps (*i.e.* 1000 random draws). For each draw we repeated our climate-matrching models testing whether (on average) birds select wintering and breeding locations that share more similar monthly precipitation values relative to null sites of the same migration distance. Out of 1000 runs, 100% had a climate matching index >0 (B). Within this randomization framework, we also tested the finding that bird in drier locations tend to exhibit more climate matching than birds in wetter locations. There was a negative effect of winter precipitation on climate matching index (based on model predicted slope) and it was significant in 92% of runs (A). Solid lines show the 'expected' value used in the formal analysis of climate matching. Dotted lines at zero represent no climate matching (B) or no effect of precipitation on climate matching (A).

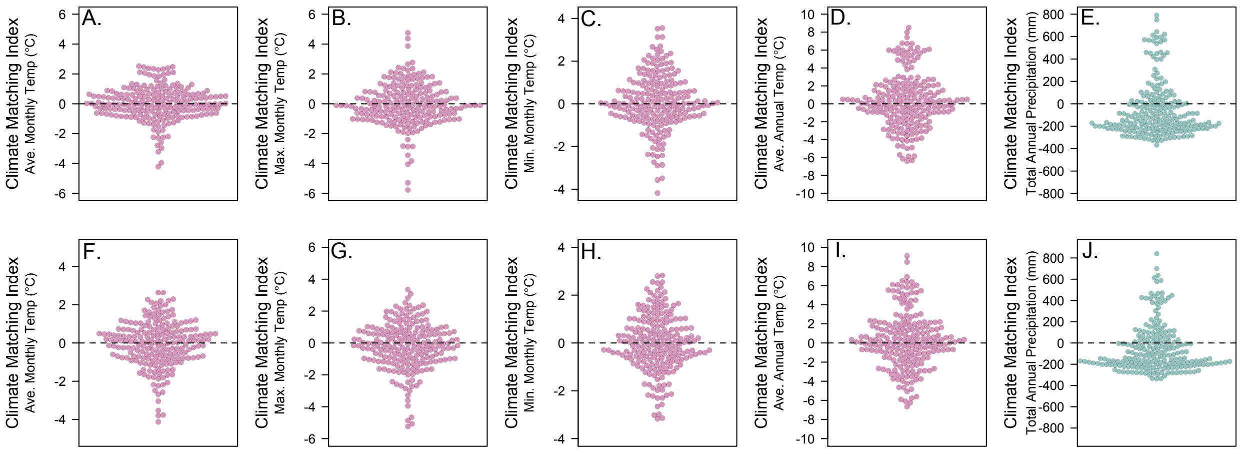


**Figure S16.** Climate matching index (CMI) based on variables extracted from the CRU (A-E) and Worldclim (F-J) dataset. Here we show the CMI for birds caught on wintering grounds, with climate represented as average monthly temperature (A,F), maximum monthly temperature (B,G), minimum monthly temperature (C,H), average annual temperature (D,I), and total annual precipitation (E,J). Points represent individuals; pink corresponds to temperature variables and blue to precipitation variables. We found no evidence of individual climate matching (CMI > 0) for any climate variable except monthly precipitation (Figures 2 and S14).

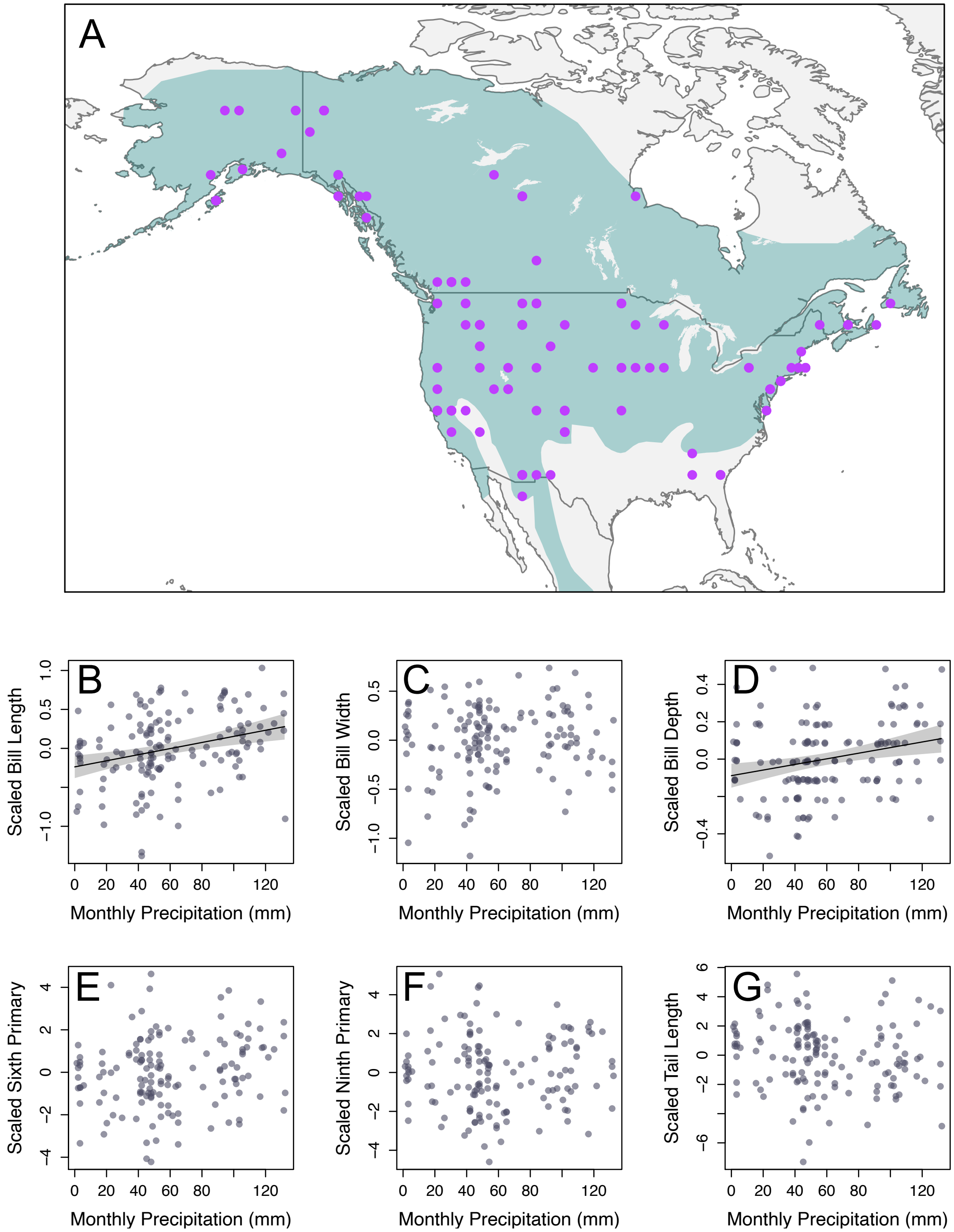


**Figure S17.** Associations between morphological data from Weidenfeld (1991) and monthly precipitation during the breeding season, measured from the CRU dataset. In the map of sample locations (A) note that locations were rounded to the nearest degree in the original dataset. Plots (B-G) show each scaled morphological measurement. Trendlines are shown in cases where a generalized least squares model was significant ($p<0.05$) after false discovery rate adjustment. Shaded regions are 95% confidence limits.

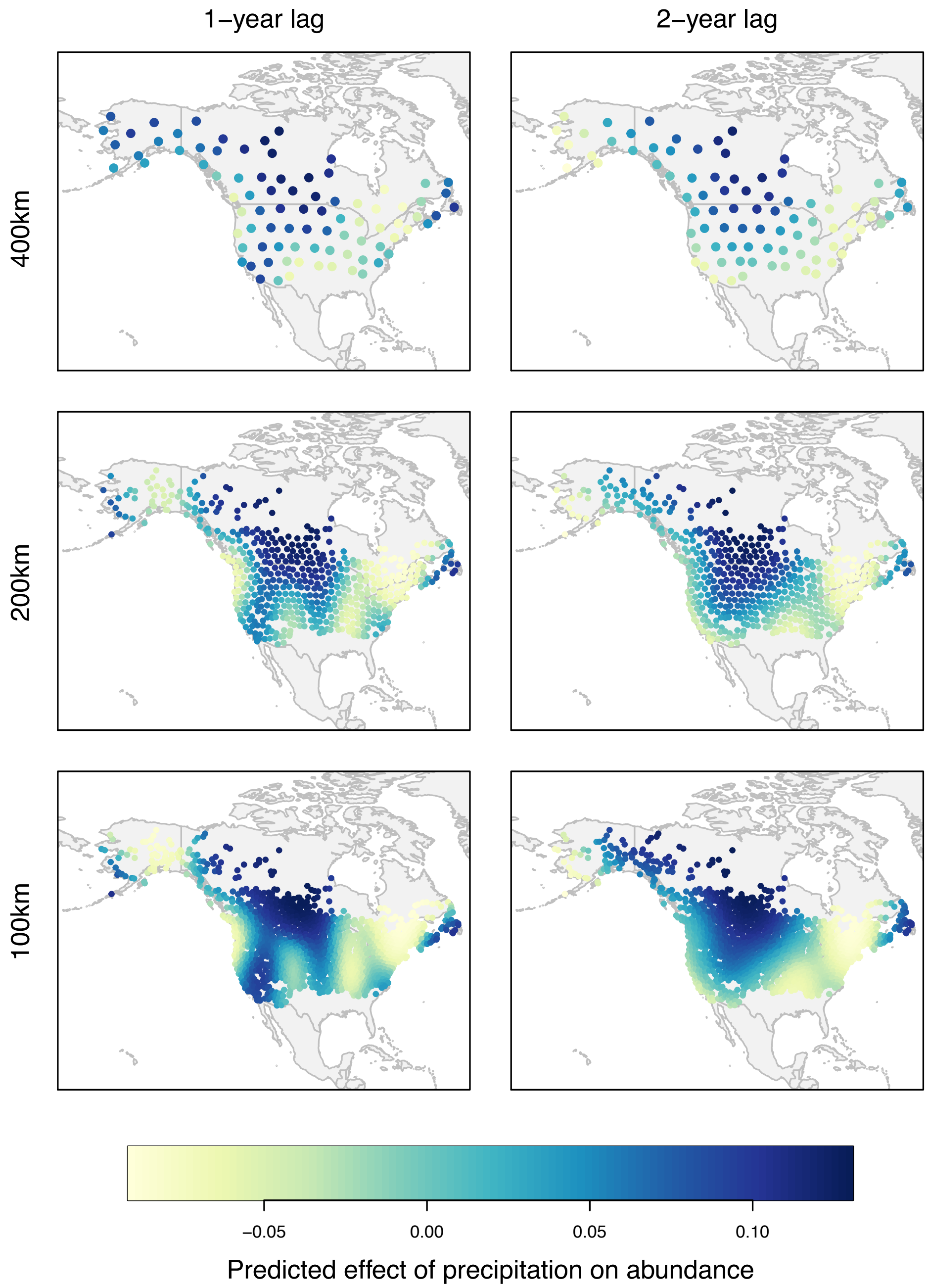


**Figure S18.** The relationship between population size and precipitation fluctuations based on three different grid cell sizes (400km, 200km, 100km). Each point is the center of a grid cell, colored by the effect of precipitation on abundance (estimated from the hierarchical model) predicted by a GAM with a spatial spline. Left panels show associations with climate for the calendar year prior to the survey and right panels show the association with climate two years prior.

**Table S1.** Sampling locations for *S. petechia* individuals sequenced for this study. For each location, we show the number of samples (n) that passed quality filters and thus were included in analysis.

| Site | State/ Province | Country | Stage | Longitude | Latitude | n | Platform |
|---|---|---|---|---|---|---|---|
| Lac La Biche | AB | USA | Breeding | -112.05 | 54.83 | 10 | RAD |
| Fairbanks | AK | USA | Breeding | -147.72 | 64.84 | 10 | RAD |
| Juneau | AK | USA | Breeding | -133.91 | 58.20 | 7 | RAD |
| Tatlayoko lake | BC | USA | Breeding | -125.42 | 51.55 | 2 | RAD |
| Fletcher lake | BC | USA | Breeding | -123.00 | 51.67 | 9 | RAD |
| Burnaby | BC | USA | Breeding | -122.70 | 49.25 | 7 | RAD |
| Revelstokes | BC | USA | Breeding | -118.20 | 51.00 | 9 | RAD |
| Creston | BC | USA | Breeding | -116.50 | 49.10 | 10 | RAD |
| Willow Creek | ID | USA | Breeding | -111.99 | 43.55 | 3 | RAD |
| Delta Marsh | MB | USA | Breeding | -98.20 | 50.20 | 9 | RAD |
| Churchill | MB | USA | Breeding | -94.12 | 58.73 | 10 | RAD |
| Germantown | NB | USA | Breeding | -64.80 | 45.68 | 4 | RAD |
| Gros Morne | NF | USA | Breeding | -57.74 | 49.69 | 10 | RAD |
| Bon Portage Island | NS | USA | Breeding | -65.70 | 43.50 | 10 | RAD |
| Queen's University Biology Station | ON | USA | Breeding | -76.32 | 44.58 | 1 | RAD |
| Chiloquin | OR | USA | Breeding | -121.93 | 42.59 | 10 | RAD |
| Trois-Rivières | QC | USA | Breeding | -88.93 | 48.30 | 10 | RAD |
| Last Mountain lake | SK | USA | Breeding | -105.23 | 51.08 | 10 | RAD |
| Moose Provincial Park | SK | USA | Breeding | -102.42 | 49.81 | 8 | RAD |
| King | WA | USA | Breeding | -122.12 | 47.56 | 6 | RAD |
| Okanogan | WA | USA | Breeding | -119.58 | 43.38 | 8 | RAD |
| Jasper | AB | USA | Breeding | -118.10 | 53.00 | 9 | Fluidigm |
| Kotzebue | AK | USA | Breeding | -162.50 | 66.90 | 10 | Fluidigm |
| Beaver Creek | AK | USA | Breeding | -141.30 | 62.70 | 2 | Fluidigm |
| Cibola | AZ | USA | Breeding | -114.70 | 33.40 | 8 | Fluidigm |
| Arcata | CA | USA | Breeding | -124.10 | 40.50 | 2 | Fluidigm |
| Alturas | CA | USA | Breeding | -120.60 | 41.50 | 9 | Fluidigm |
| San Luis Obispo | CA | USA | Breeding | -120.50 | 35.20 | 1 | Fluidigm |
| Lee Vining | CA | USA | Breeding | -119.10 | 37.90 | 8 | Fluidigm |
| Kent | CT | USA | Breeding | -74.36 | 40.70 | 6 | Fluidigm |

| | | | | | | | |
|---|---|---|---|---|---|---|---|
| Junction City | KS | USA | Breeding | -96.83 | 39.10 | 12 | Fluidigm |
| Wells | ME | USA | Breeding | -70.55 | 43.30 | 11 | Fluidigm |
| Vicksburg | MI | USA | Breeding | -85.52 | 42.20 | 11 | Fluidigm |
| FCTC-SABO | MI | USA | Breeding | -84.70 | 42.30 | 10 | RAD |
| Dearborn | MI | USA | Breeding | -83.24 | 42.30 | 6 | Fluidigm |
| North Oaks | MN | USA | Breeding | -93.06 | 45.10 | 11 | Fluidigm |
| Ravalli | MT | USA | Breeding | -114.20 | 47.30 | 10 | Fluidigm |
| Holter Dam | MT | USA | Breeding | -111.90 | 46.90 | 9 | Fluidigm |
| Denton | MT | USA | Breeding | -109.70 | 47.40 | 3 | Fluidigm |
| Kent Island | NB | USA | Breeding | -66.80 | 44.60 | 7 | Fluidigm |
| Layton | NJ | USA | Breeding | -74.40 | 41.10 | 5 | Fluidigm |
| Ruby Valley | NV | USA | Breeding | -115.50 | 40.20 | 10 | Fluidigm |
| Brockport | NY | USA | Breeding | -77.88 | 43.20 | 8 | Fluidigm |
| Stanfordville | NY | USA | Breeding | -73.68 | 41.90 | 11 | Fluidigm |
| Garfield Hts. | OH | USA | Breeding | -81.65 | 41.40 | 9 | Fluidigm |
| Klamath | OR | USA | Breeding | -121.92 | 42.68 | 9 | Fluidigm |
| Bethlehem | PA | USA | Breeding | -75.38 | 40.62 | 20 | RAD |
| Montreal | QC | USA | Breeding | -73.52 | 45.70 | 8 | Fluidigm |
| Midway | UT | USA | Breeding | -111.50 | 40.50 | 12 | Fluidigm |
| Silverton | WA | USA | Breeding | -121.41 | 48.08 | 5 | Fluidigm |
| Goose Prairie | WA | USA | Breeding | -121.30 | 47.00 | 12 | Fluidigm |
| Ferry | WA | USA | Breeding | -118.60 | 48.54 | 1 | Fluidigm |
| San Miguelito | Ahuachapan | El Salvador | Wintering | -89.94 | 13.80 | 1 | Fluidigm |
| San José del Cabo | Baja California Sur | Mexico | Wintering | -109.68 | 23.05 | 2 | Fluidigm |
| Granada | Granada | Nicaragua | Wintering | -86.01 | 11.80 | 1 | Fluidigm |
| Tamarindo | Guanacaste | Costa Rica | Wintering | -85.83 | 10.32 | 18 | Fluidigm |
| Liberia | Guanacaste | Costa Rica | Wintering | -85.67 | 10.78 | 17 | Fluidigm |
| Guardia | Guanacaste | Costa Rica | Wintering | -85.62 | 10.60 | 27 | Fluidigm |
| Chamela | Jalisco | Mexico | Wintering | -105.08 | 19.53 | 12 | Fluidigm |
| Turrialba | Jinotega | Nicaragua | Wintering | -86.05 | 13.20 | 18 | Fluidigm |
| El Quince | Merida | Venezuela | Wintering | -71.77 | 8.58 | 21 | Fluidigm |
| West Michoacan | Mitchoacan | Mexico | Wintering | -101.11 | 19.70 | 14 | Fluidigm |
| Manialtpec | Oaxaca | Mexico | Wintering | -96.75 | 17.08 | 1 | Fluidigm |
| Chila | Oaxaca | Mexico | Wintering | -96.65 | 15.95 | 3 | Fluidigm |
| Hidalgo | Oaxaca | Mexico | Wintering | -96.65 | 15.95 | 2 | Fluidigm |
| Oaxaca | Oaxaca | Mexico | Wintering | -96.65 | 15.95 | 1 | Fluidigm |

| | | | | | | | |
|---|---|---|---|---|---|---|---|
| Paisley | San Jose | Costa Rica | Wintering | -82.95 | 8.75 | 10 | Fluidigm |
| Santa Teresa | San Jose | Costa Rica | Wintering | -82.92 | 8.81 | 20 | Fluidigm |
| Metapan | Santa Ana | El Salvador | Wintering | -89.36 | 14.40 | 4 | Fluidigm |
| Izalco | Sonsonate | El Salvador | Wintering | -89.65 | 13.80 | 9 | Fluidigm |
| Salinas | Veracruz | Mexico | Wintering | -95.17 | 18.50 | 18 | Fluidigm |
| Celestun | Yucatan | Mexico | Wintering | -90.40 | 20.87 | 4 | Fluidigm |

**Table S2.** Summary of pre- and post-filtering sample sizes and number of loci for each genotyping method.

| Method | Pre-filtering SNPs | Filtered SNPs | Pre-filtering N | Filtered N | Stage |
|---|---|---|---|---|---|
| RAD | 4,335,072 | 104,711 | 229 | 195 | Breeding |
| Fluidigm | 192 | 157 | 231 | 224 | Breeding |
| Fluidigm | 96 | 96 | 203 | 203 | Wintering |

**Table S3.** Evidence for individual-level niche tracking for monthly precipitation but not temperature or annual climatic conditions (*i.e.,* temperature and precipitation calculated across the entire year, even when the species is absent). First number depicts the mean predicted Climate Matching Index; parentheses depict 95% confidence intervals. Significant effects are bolded (*P*< 0.05). Analyses using both CRU and WorldClim climate data are shown.

| | Average Monthly Precipitation | Average Monthly Temperature | Maximum Monthly Temperature | Minimum Monthly Temperature | Average Annual Temperature | Total Annual Precipitation |
|---|---|---|---|---|---|---|
| CRU | **15.0 (9.1, 20.9)** | -0.02 (-0.34, 0.30) | -0.08 (-0.47, 0.30) | 0.03 (-0.44, 0.50) | 0.28 (-0.58, 1.14) | -50.4 (-115.5, 15.5) |
| WorldClim | **12.9 (6.8, 19.0)** | -0.14 (-0.45, 0.16) | -0.26 (-0.60, 0.07) | -0.14 (-0.54, 0.25) | 0.23 (-0.60, 1.07) | -40.7 (-101.5, 21.5) |

**Table S4.** Analyses of individual-level niche tracking with respect to monthly precipitation. Table shows results from two types of Linear Mixed Models (LMMs). The first estimates how interactions between winter precipitation and the type of prediction (*i.e.,* actual predicted breeding locations from sequence data vs. null predictions only accounting for distance) affect predicted breeding locations. The second type of LMM assesses effects of winter precipitation on the climate matching index. Quadratic and linear effects are included. Columns depict coefficient estimates from LMMs, and $\chi2$ and *P*-values comparing nested models via Likelihood Ratio Tests.

| | Quadratic interactive effect of winter precipitation and type of prediction (actual prediction vs. null; *i.e.,* Fig. 2a) | | | Linear interactive effect of winter precipitation and type of prediction (actual prediction vs. null; *i.e.,* Fig. 2a) | | | Quadratic effect of winter precipitation on climate matching index (*i.e.,* Fig. 2b) | | | Linear effect of winter precipitation on climate matching index (*i.e.,* Fig. 2b) | | |
|---|---|---|---|---|---|---|---|---|---|---|---|---|
| | Coefficient | $\chi2$ | *P*-value | Coefficient | $\chi2$ | *P*-value | Coefficient | $\chi2$ | *P*-value | Coefficient | $\chi2$ | *P*-value |
| CRU | -8.13 | 28.4 | <0.001 | 18.24 | 92.4 | <0.001 | 6.43 | 4 | 0.041 | -12.2 | 10.1 | 0.001 |
| WorldClim | -4.66 | 3.88 | 0.049 | 12.52 | 40 | <0.001 | Did not converge | | | -7.34 | 7 | 0.008 |
